# Investigating Invariances in Auditory Event Categorization with Model Metamers

**DOI:** 10.64898/2026.07.31.741881

**Authors:** Hee So Kim, Elizabeth Lee, Malinda McPherson-McNato, Abigail Noyce, Jenelle Feather

## Abstract

Real-world acoustic inputs contain rich sensory information that we parse into discrete auditory objects and categories. Although deep neural networks (DNNs) are increasingly used to model auditory perception, the field lacks rigorous behavioral benchmarks, particularly for auditory event categorization. Here, we developed a 25-way categorization paradigm for broad classes of natural sounds to test whether categorical invariances of DNNs align with those of human observers. We first confirmed that humans could reliably categorize the natural sounds, demonstrating that our paradigm is well-suited for testing invariances in auditory categories. To probe model invariances, we evaluated human recognition of ‘model metamers’ (synthetic stimuli matched to the model’s internal activations for each natural stimulus) for a wide range of architectures trained on speech or auditory event recognition. Evaluating widely-used public models, we found that human recognition of auditory event model metamers was generally influenced by the training task and data distribution; speech models trained on standard, curated datasets produced less recognizable metamers than auditory event recognition models. We additionally analyzed a controlled set of models to directly investigate the influence of training task and adversarial training, revealing that improved metamer recognition induced by adversarial training is task-dependent. However, even in the best-performing models, we observed a sharp decline in human recognition at the final classification layer compared to the penultimate representation layer. Overall, our results suggest that while invariances in modern architectures better align with human observers for auditory event categorization, there is still a large discrepancy between the categorical invariances of auditory neural networks and the invariances of human observers. Code and models are available at https://github.com/Feather-Lab/env-sound-metamers

## Introduction

A central challenge of auditory perception is mapping highly variable acoustic waveforms into stable, semantic categories. For example, we can easily recognize the sound of a dog barking despite vast acoustic differences between the deep bark of a Mastiff or the yip of a Chihuahua. While this categorical invariance is critical for perception (DiCarlo & Cox, 2007), the computational mechanisms underlying how the human auditory system generalizes across highly varying acoustic features to form distinct categories remain poorly understood (Banno et al., 2020). Historically, most studies of auditory categorization have focused on domains of speech (Mann, 1980; Holt, 2006), music (Locke and Kellar, 1973; Perrot, 1999), or synthetic tone sequences (Ohl et al., 2001). In these domains, stimuli often possess structured parametric features (e.g., phonemes or pitch changes) that allow researchers to systematically investigate categorization by manipulating the acoustic attributes. However, because many natural sounds cannot be easily manipulated parametrically, it is difficult to scale some of these approaches to more varied auditory contexts. Thus, these studies may be missing critical aspects of the cognitive translation from acoustic inputs into semantic, categorical representations of natural sounds.

Task-optimized deep neural networks (DNNs) offer a powerful framework to investigate these complex auditory computations. Because auditory DNNs transform raw waveforms into high-level representations, they offer an opportunity to explore categorical invariances for naturalistic auditory events. We can consider this type of computational model as an instantiation of a hypothesis about human representations, but such a model must be critically evaluated for alignment or misalignment with human perception. Evaluating the “alignment” between deep neural networks and human perception is a complex challenge, as the term encompasses a broad taxonomy of behavioral, structural, and representational similarities, and model results can vary depending on the precise domain being investigated (Sucholutsky* et al., 2023). While DNN representations have been widely compared to human representations in domains such as speech recognition (Adolfi et al., 2023; Kell et al., 2018; Li et al., 2023; Millet et al., 2022), music classification (Güçlü et al., 2016; Kell et al., 2018), and pitch perception (Saddler et al., 2021), systematic evaluations of model-human similarities for broad classes of natural sounds remain limited (Alavilli & McDermott, 2026). Some work has focused on using auditory-event trained DNNs to predict fMRI activity for natural sounds (Giordano et al., 2023; Tuckute et al., 2023), however this work has focused on fairly small dataset sizes, and neural prediction does not directly test behavioral alignment. Other lines of work have directly compared human and model behavior, but this has either been restricted to narrow domains such as impact sounds (Bost et al., 2024), focused on continuous perceptual similarity rather than explicit categorization (Giordano et al., 2023), or based on target detection of a probed category in an auditory scene (Alavilli & McDermott, 2026). Consequently, it remains an open question how well modern DNNs align with human behavior when categorizing a diverse set of everyday auditory events. In this work, we focus on comparing the categorical invariances of deep neural networks to the invariances of human observers. We develop a novel 25-way forced-choice categorization task to serve as a benchmark for model-human behavioral alignment across auditory event categories. We validate this task with natural auditory events, and then use the task to compare human and model invariances via the framework of model metamers (Feather et al., 2019, 2023). Model metamers are two stimuli that are physically distinct but produce the same activation pattern within a computational model. In the context of a classification task, we can take a natural stimulus (for instance, the sound of a dog) and find a new sound that generates the same activation pattern in a model, leading the model to predict with equal confidence that the new sound is the same class as the original sound. Thus, the overall motivation of this work is twofold: first, to introduce a rigorous behavioral benchmark for auditory event categorization, and second, to utilize this benchmark to quantify auditory categorization differences between human perception and modern deep neural networks. Specifically, our main contributions are:

1. We develop an engaging 25-way forced-choice auditory event categorization task that enables direct quantitative comparisons between the behavior of DNNs and human observers.
2. We generate model metamers for auditory events from DNNs optimized for speech recognition and auditory event recognition to investigate the task-specificity of learned representational invariances.
3. Consistent with previous findings in word recognition and image classification, we demonstrate that adversarial training significantly improves alignment of invariances between DNNs and human observers for auditory event categories.

## Auditory Event Categorization Task

The auditory domain currently lacks a standardized framework for comparing DNN performance to human perception for broad categories of natural sounds. Engineering-focused models are frequently evaluated on large-scale datasets that can present a challenge for direct comparison with human observers. For example, AudioSet (Gemmeke et al., 2017) contains 527 hierarchically organized labels, which present challenges due to its massive, multi-label ontology where a single clip simultaneously belongs to many classes (i.e. a sound is labeled as music, musical instrument, plucked string instrument, and banjo), making it ill-suited for a forced-choice categorization task. In the auditory perception domain, existing human behavioral tasks often rely on paradigms such as free-sorting tasks (Collett et al., 2016, Berland et al., 2019) that are difficult to implement or evaluate in models optimized for classification.

To address this, we drew inspiration from work in visual object recognition, which utilized a 16-class image classification task to create a common ground for comparison between models and humans (Geirhos et al., 2018). We developed a 25-way forced-choice sound categorization task using a gamified icon-selection paradigm (Figure 1a; icons from The Noun Project). This constrained forced-choice paradigm is advantageous because it avoids the high response variability of open-ended identification tasks, where human synonyms are difficult to reconcile with a model’s fixed vocabulary. In addition, using icons on the response screen instead of text makes the task accessible and engaging, encouraging high quality behavioral data.

**Figure 1:**
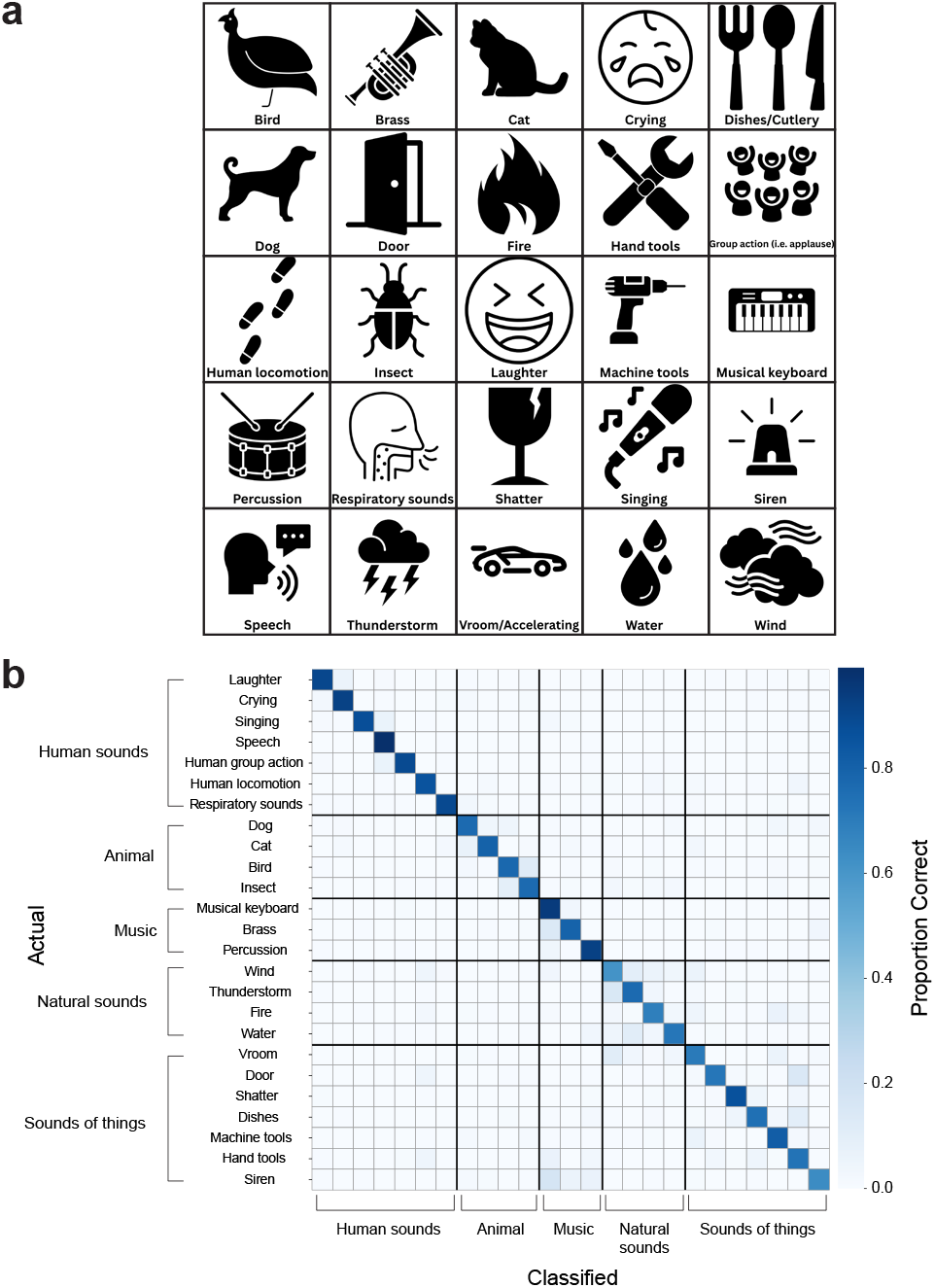
In all experiments, listeners heard stimuli sequentially and assigned each to one of 25 categories. (a)Participants were instructed to click one category from the response display comprising a 5×5 grid with an icon and label for each category option. (b) Confusion matrix for human observers (*N* = 20) from the natural sounds experiment, with actual sound categories in rows and human responses in columns. Human responses are overwhelmingly along the main diagonal, indicating high classification accuracy. Average task accuracy = 0.804 *±* 0.019 (SEM across participants).

### Category and stimuli selection

To select our 25 categories, we chose labels from the AudioSet ontology (Gemmeke et al., 2017) that represent common and salient environmental sounds. We intentionally selected categories from various “heights” of the taxonomic tree to capture a range of semantic breadths. For example, our set includes broad categories like “water” as well as more specific, narrow classes such as “hand tools.” This diversity ensures the benchmark captures a wide range of semantic granularities found in auditory scenes.

To generate an interpretable set of environmental sound recordings for evaluation, we selected audio clips from the evaluation set of the FSD50K dataset (Fonseca et al., 2022). We applied a strict filtering procedure to ensure semantic clarity, including only those clips that possessed a single annotated category label from our target set. Using the hierarchical structure of the AudioSet ontology, we resolved parent-child relationships to ensure that sounds were not flagged as multi-labeled simply because they possessed both a specific label (e.g., “Hammer”) and its broader parent class (e.g., “Tools”). However, we strictly excluded any clips that contained multiple distinct sources from our target set, such as a recording featuring both a “hand tool” and a “machine tool.” This rigorous mapping ensured that each of the 275 stimuli uniquely represented a single category, minimizing ambiguity for both human listeners and model evaluations. The final stimulus set included 11 unique sound clips per category, resulting in a total of 275 stimuli, each 2 seconds in duration.

We chose FSD50K as our source evaluation set for three primary reasons. First, FSD50K utilizes the AudioSet ontology, allowing standard evaluation of models that are pretrained on AudioSet without requiring posthoc interventions to reconcile distinct label sets, for instance via fine-tuning a model or training a linear classifier (as typically done for Piczak (2015)). Second, as several of these models were optimized or selected based on their AudioSet performance, we did not want to “double dip” and assess them on that same corpus. Finally, FSD50K provides more rigorously annotated labels than the original AudioSet, ensuring a reliable ground-truth for our human behavioral comparisons.

### Human Behavioral Experiment: Natural Sounds

Behavioral experiments for the natural sound categorization task were conducted on Prolific (full details in SM.2). All participants completed 275 experimental trials, where each trial consisted of a single presentation of a unique two-second sound clip, and participants chose the category to which they believed the sound belonged by clicking on the icon corresponding to the category. Prior to the main experiment, participants completed a familiarization and practice phase (full trial structure and training details provided in SM.2). During the main experiment, participants were only able to listen to each two-second sound clip once. Following the click of the icon, there was a 250ms inter-trial interval before the next sound was played.

## Results: Human categorization of natural sounds

To establish a behavioral baseline for auditory event recognition, we evaluated human performance on the 275 natural sound recordings. Human observers (*N* = 20) achieved an average accuracy of 80.4% ± 1.9% (SEM across participants). As shown in the confusion matrix in Figure 1b, human responses were concentrated heavily along the main diagonal. This high level of accuracy and the clear diagonal structure validate our experimental design, confirming that the 25-way icon-matching task is an effective paradigm for capturing human category knowledge. Additionally, the confusion patterns themselves were fairly reliable across participants (split-half reliability = 0.706, Spearman-Brown corrected Spearman correlation, off-diagonal cells only), suggesting that some of the sounds may be semantically ambiguous when presented in isolation. Overall, these results demonstrate that our curated stimulus set is highly and reliably recognizable to human listeners and that the 25-way classification task provides a robust benchmark for comparing human and model representations.

## Human-model alignment on auditory event categorization

To establish a baseline for human-model alignment, we evaluated a suite of models explicitly optimized for auditory event categorization on the 25-way task.

### Auditory Event Models

We evaluated five models trained on auditory event recognition (Table S2). These models were selected to provide a representative sample of modern architectures that span different model classes and training procedures. We tested three publicly available state-of-the-art models trained on audio event categorization: AST (Gong et al., 2021), BEATs (Chen et al., 2022), and Whisper-AT (Gong et al., 2023) (summary of key model components, architectural and training details for all models are provided in Table S1). We additionally tested two CochResNet50 models that were trained on the Word-Speaker-Noise dataset (Feather et al., 2019). Each model consisted of a biologically inspired “cochleagram” front-end (Glasberg and Moore, 1990; McDermott and Simoncelli, 2011) followed by a ResNet50 (He et al., 2016) architecture. We analyzed a model pretrained on the Audio Event Classification component of the Word-Speaker-Noise (WSN) task (CochEvent-S; the CochResNet50-AudioSet from Tuckute et al., 2023). We additionally trained a model using the same architecture, dataset, and Audio Event Classification task as reported in (Tuckute et al., 2023), but trained to be resilient to cochleagram-based adversarial attacks (Feather et al., 2023; Madry et al., 2017). Adversarial examples are small, often human-imperceptible perturbations that are designed to change the model’s classification decision (Goodfellow et al., 2014; Szegedy et al., 2013). Adversarial training adds small perturbations to each training example during the optimization process to explicitly force the model to classify these examples correctly (Madry et al., 2017). Previous work has demonstrated that this process can improve the recognizability of metamers for image classification and word recognition (Feather et al., 2023), even if it slightly reduces the overall classification accuracy (Tsipras et al., 2018). Training details for the CochEvent-R model are given in Supplemental Material SM.1.

To allow direct comparison between model and human behavior, we implemented a 25-way forced-choice categorization task for all models by aggregating model predictions across all sub-categories relevant to our 25 classes and requiring the model to select the category with the highest mean prediction, following previous work (Geirhos et al., 2018). All pretrained models were obtained using the HuggingFace Transformers library (Wolf et al., 2019) or their respective official GitHub repositories.

## Results: Model performance on 25-way categorization task

For models trained on auditory event categorization, we evaluated how well the models categorized natural sound stimuli compared to human performance. All models performed significantly above the chance level of 0.04, though overall accuracy varied considerably across architectures (Figure 2a). Among the audio event models, AST achieved the highest performance (77.2% ± 2.5%), closely approaching the human baseline of 80.4%. In contrast, other models such as CochEvent-S (69.1%) and Whisper-AT (68.4%) showed moderate performance, while CochEvent-R (52.4%) and BEATs (49.8%) exhibited notably lower accuracy on this 25-way task. To determine whether models and humans share similar strategies, we compared their error patterns by calculating the Spearman correlation between their respective confusion matrices (Figure 2b & c). We analyzed the confusion matrix both including and excluding the diagonal, to capture both correct classifications and specific patterns of mis-categorization. To establish human-human error consistency, we calculated the mean human splithalf reliability (Spearman correlation over 1,000 random participant splits), which then was Spearman-Brown corrected. Although the models showed moderate positive correlations with human behavior, these correlations were significantly lower than the human split-half reliability. This gap indicates that even when models achieve high accuracy on the categorization task, the specific errors they make differ from those made by human observers. This was especially notable when comparing only the incorrect trials (the off-diagonal elements of the confusion matrix). These results suggest that standard training objectives and architectures may not yet fully capture the representations used by the human auditory system for categorization tasks.

**Figure 2:**
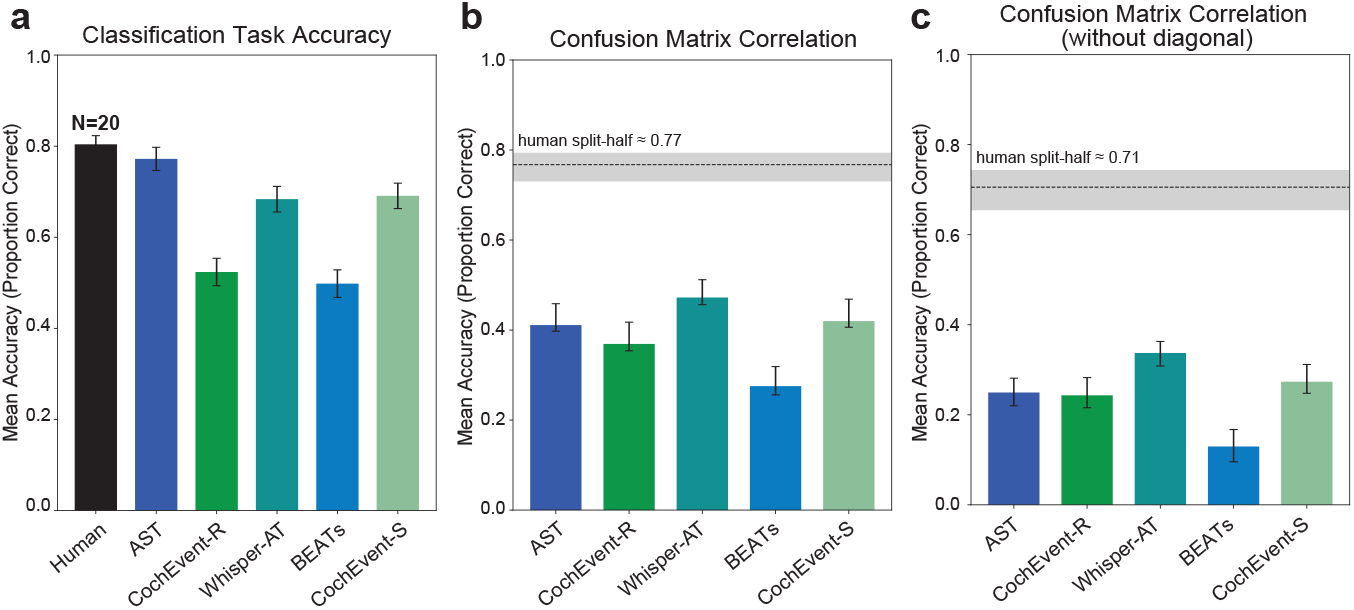
(a) Human and auditory event model categorization accuracy on the 25way task. Error bars denote SEM across sounds. (b & c) Correlations between the confusion matrix (b: including diagonal; c: excluding diagonal) for human observers and each model are moderate and positive, but lower than human split-half reliability (black dashed line). Shaded line and error bars denote 95% confidence intervals (CIs). For models, CIs were obtained by bootstrapping participants (1,000 resamples).

## Investigating invariance alignment using model metamers

While our previous analysis of error patterns suggests a divergence between humans and models, alternative approaches utilize synthetic stimuli optimized to reveal properties of the representations of a computational model (Berardino et al., 2017; Feather et al., 2024; Golan et al., 2020; Wang & Simoncelli, 2008). Here, we use the framework of model metamers: two stimuli that are physically distinct but that produce the same activation patterns within a computational model. Model metamers can be utilized to probe the internal invariances of a neural network, as the stimuli differ in ways that a model is invariant to, making them well suited to study aspects of categorical perception. Prior work on model metamers has primarily explored the domains of word and visual object recognition (Feather et al., 2019, 2023). In both cases, metamers generated from late model stages often become completely unrecognizable to human observers, suggesting a significant divergence between model and human invariances. We were motivated by these findings to apply this framework to the understudied domain of auditory event recognition and to investigate the emergence of task specific invariances and the benefits of adversarially robust representations.

### Model metamer generation

We generated model metamers using an iterative optimization procedure detailed in Feather et al., 2023. For each metamer, we initialized a signal with noise and used gradient descent to minimize the normalized *L*_2_ distance between the activations of this synthetic signal and those of the paired natural signal (Figure 3a). All model weights remained fixed during this process. To ensure stable optimization, we applied gradient clipping to operations that typically cause numerical instability, such as logarithms and power functions common in audio preprocessing. We also performed two validation steps to confirm that the optimization was successful. First, we calculated the Spearman correlation between metamer activations and the original signal; all models achieved correlation values above 0.988. Second, we applied a strict inclusion criterion: the classification of the synthesized metamer by the model had to match its classification of the original natural stimulus. See Supplementary Table S3 for details on the optimization parameters.

**Figure 3:**
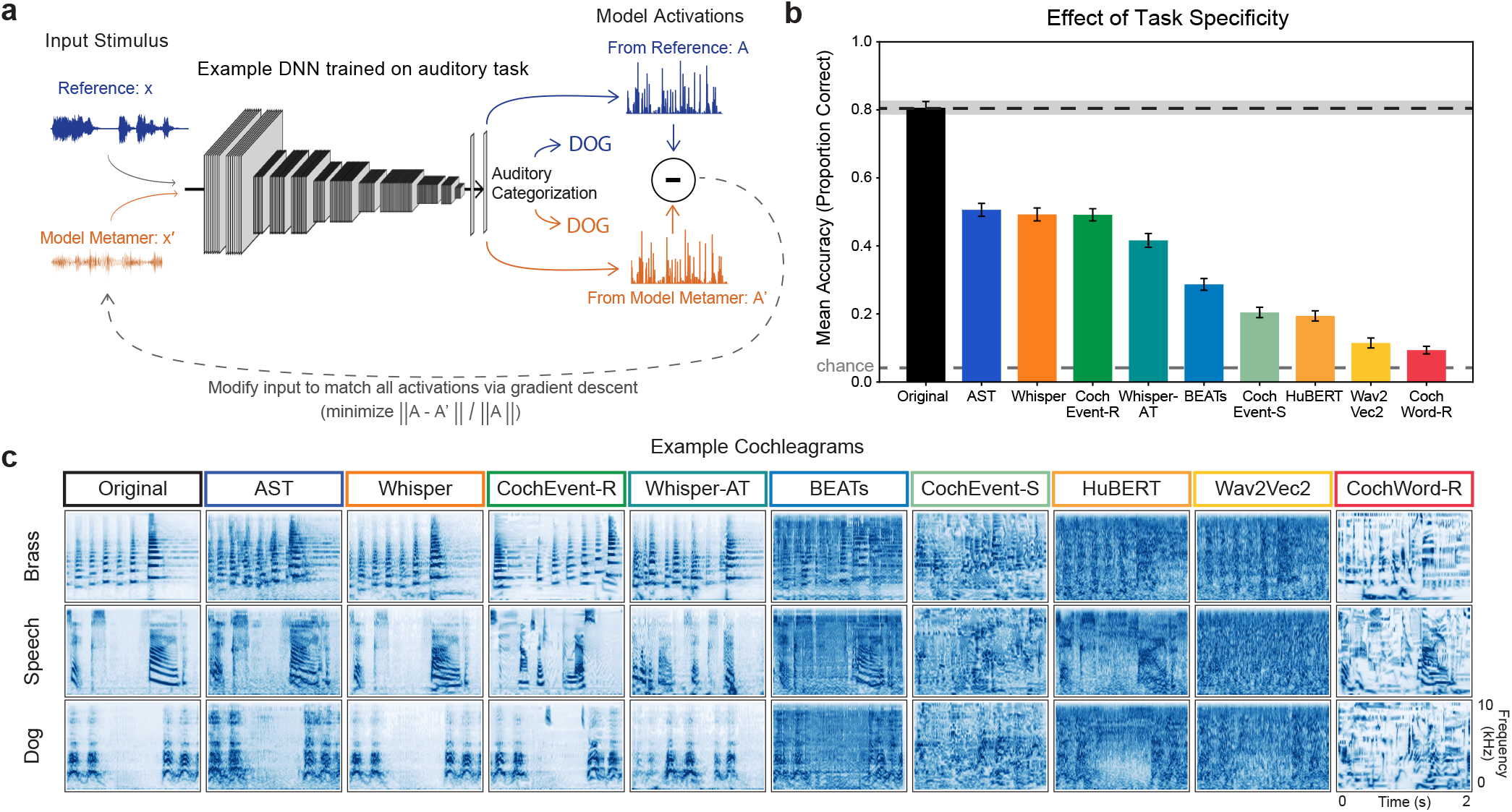
(a) Schematic of the model metamer generation process (adapted from Feather et al., 2023). Metamers were synthesized by using gradient descent to minimize the normalized *L*_2_ distance between the activations of a synthetic signal and those of a paired natural signal. (b) Human recognition of model metamers across all nine models compared to that of natural sounds (*N* = 48). Dotted line denotes human recognition accuracy of natural sounds. Shaded line and error bars denote SEM across participants. Metamers were generated from the penultimate representation layer of each model. (c) Example cochleagrams from “Brass”, “Speech”, and “Dog” categories for all nine models, shown alongside the specific natural sound used as the reference for its synthesis.

### Speech recognition models

In addition to the five models trained on audio event recognition (described above), we tested four models trained on speech recognition to evaluate how task-specificity interacts with model invariances (Table S2). We included three state-of-theart speech recognition models: Wav2Vec 2.0 (Baevski et al., 2020), HuBERT (Hsu et al., 2021), and Whisper (Radford et al., 2023). We also tested CochWord-R (Feather et al., 2023), which shares the same training dataset as well as model architecture as CochEvent-S and CochEvent-R, but was trained on the word recognition task from the Word-Speaker-Noise (WSN) dataset, and was trained with *l*_2_-norm(ϵ = 1) adversarial examples on the cochleagram representation.

### Layer selection

Following previous research showing that model invariances diverge from human perception in deeper stages (Feather et al., 2019, 2023), we targeted the penultimate “representation layer” (the layer preceding the classification head) for all nine models. For the five models trained specifically for audio event categorization (AST, BEATs, Whisper-AT, CochEvent-S, and CochEvent-R), we additionally tested metamers from the final classification layer (Table S2).

### Human Behavioral Experiment: Metamer Experiments

In this experiment, 48 participants performed the 25-way categorization task while listening to both model metamers and natural sounds. We presented 275 unique sounds across 15 different model and layer conditions. These conditions included metamers generated from the representation layers of nine different models, metamers from the final classification layers of five models trained for audio event recognition, and one condition with natural sounds. Conditions were randomized across the 275 unique sounds so that each participant completed a median of 18 trials per condition (range 16-19), and to avoid priming, no unique sound was presented in multiple conditions to the same observer. Additionally, from the total pool of 4,125 possible stimuli (representing the combination of all sounds and conditions), we ensured that each individual sound was presented at least twice (maximum four times) across the entire group of participants. The trial structure, introduction, and training phases followed the same structure as the natural sounds experiment.

## Results: Human recognition of model metamers

### Effect of Task Specificity

To investigate whether the internal representations of these models align with human category invariances, we tested how well human observers could categorize metamers generated from each model’s penultimate representation layer. While humans easily categorized the original natural sounds, their performance on metamers varied significantly across models (Figure 3).

We first examined whether the specific task a model was trained to perform influences the recognizability of its metamers. Broadly speaking, metamers generated from models trained for audio event categorization were more recognizable compared to those generated from models trained for speech recognition (Figure 3b,c). This pattern suggests that representations optimized for auditory event categorization tend to exhibit more human-like invariances for natural sounds than representations optimized for speech recognition. One significant exception to the overall low-accuracy for speech recognition models was Whisper. Despite being trained for various speech recognition tasks but not any audio event recognition tasks, Whisper produced highly recognizable metamers and achieved the second-highest accuracy across all model conditions. A closer look at individual auditory categories revealed that some other speech models (such as HuBERT and CochWord-R) had largely unrecognizable model metamers for all sound categories except speech, where recognition of the metamer as a speech stimulus was high (Figure S2). We additionally observed notable variability in the human recognizability of model metamers among the audio event categorization models, which was particularly evident when analyzing recognition accuracy individually for each auditory category. These differences likely arise from variations in architecture, dimensionality, and specific training objectives.

### Effect of Adversarial Training

To examine how adversarial training influences the recognizability of model metamers, we tested the recognition of metamers generated from three controlled models: CochEvent-S, CochEvent-R, and CochWord-R (Figure 4; results for these models are replotted from Figure 3 to facilitate direct comparison). These models share the same CochResNet50 architecture and training data, but differ in the objective functions used for training. Across all model conditions, human accuracy for model metamer recognition was lower than the recognition accuracy for natural sounds. However, the model metamers generated from CochEvent-R (the adversarially trained audio event model) were more recognizable than the model metamers generated from CochEvent-S (the standard trained audio event model). Thus, consistent with previous work in the word and image recognition domains (Feather et al., 2023), adversarial training led to a reliable improvement in the perceptual recognizability of metamers. CochWord-R (the adversarially trained word recognition model) showed the lowest recognizability among the three model conditions, even though this model was previously reported to have highly recognizable model metamers in the domain of word recognition (Feather et al., 2023). This dissociation shows that adversarial training alone is not sufficient to improve metamer recognizability, and rather the task on which the model is trained plays a critical role in shaping the model invariances.

**Figure 4:**
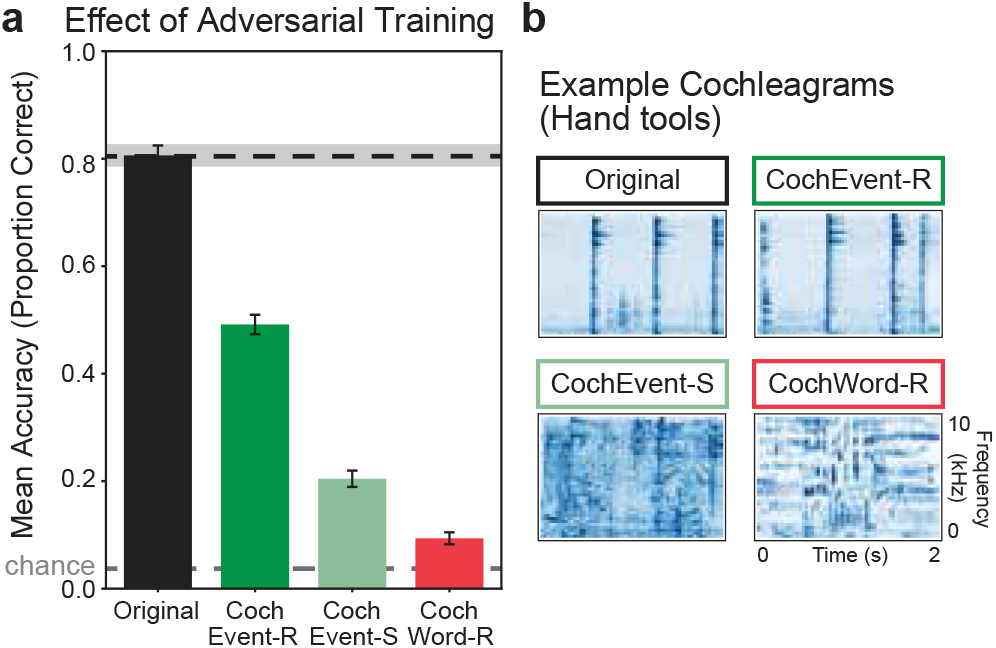
(a) Human recognition of model metamers across three CochResNet50 models compared to that of natural sounds (*N* = 48). Dotted line denotes human recognition accuracy of natural sounds. Shaded line and error bars denote SEM across participants. Results for these models are replotted from Figure 3 to facilitate direct comparison. (b) Example cochleagrams of a reference natural sound from the “Hand tools” category and corresponding model metamers.

### Effect of Metamer Generation Stage

Lastly, for models trained on audio event recognition, we examined whether the stage of metamer generation influenced recognizability. Specifically, we compared metamers generated from two model stages: the penultimate representation layer and the final classification layer. We tested metamers from the final classification layer because it is the stage at which category decisions are computed and where task-relevant categorical invariances are most directly reflected.

Across models, metamers generated from the final classification layer were less recognizable than those generated from the representation layer. However, among these models, the adversarially trained model produced the most recognizable metamers from the final classification layer (Figure 5). To better understand where the divergence occurs between the standard and adversarially trained models across the model hierarchy, we conducted an additional behavioral experiment testing intermediate stages of the CochResNet50 architectures (Figure 6). We observed that human recognition of metamers remains high in early layers, declines gradually in middle stages, and drops sharply at the final task-specific classification stage. Notably, this decrease is less pronounced with adversarially trained models. Although adversarial training appears to make model invariances more aligned with that of humans, the overall pattern indicates that the invariances supporting model category decisions still differ substantially from those used by human listeners.

**Figure 5:**
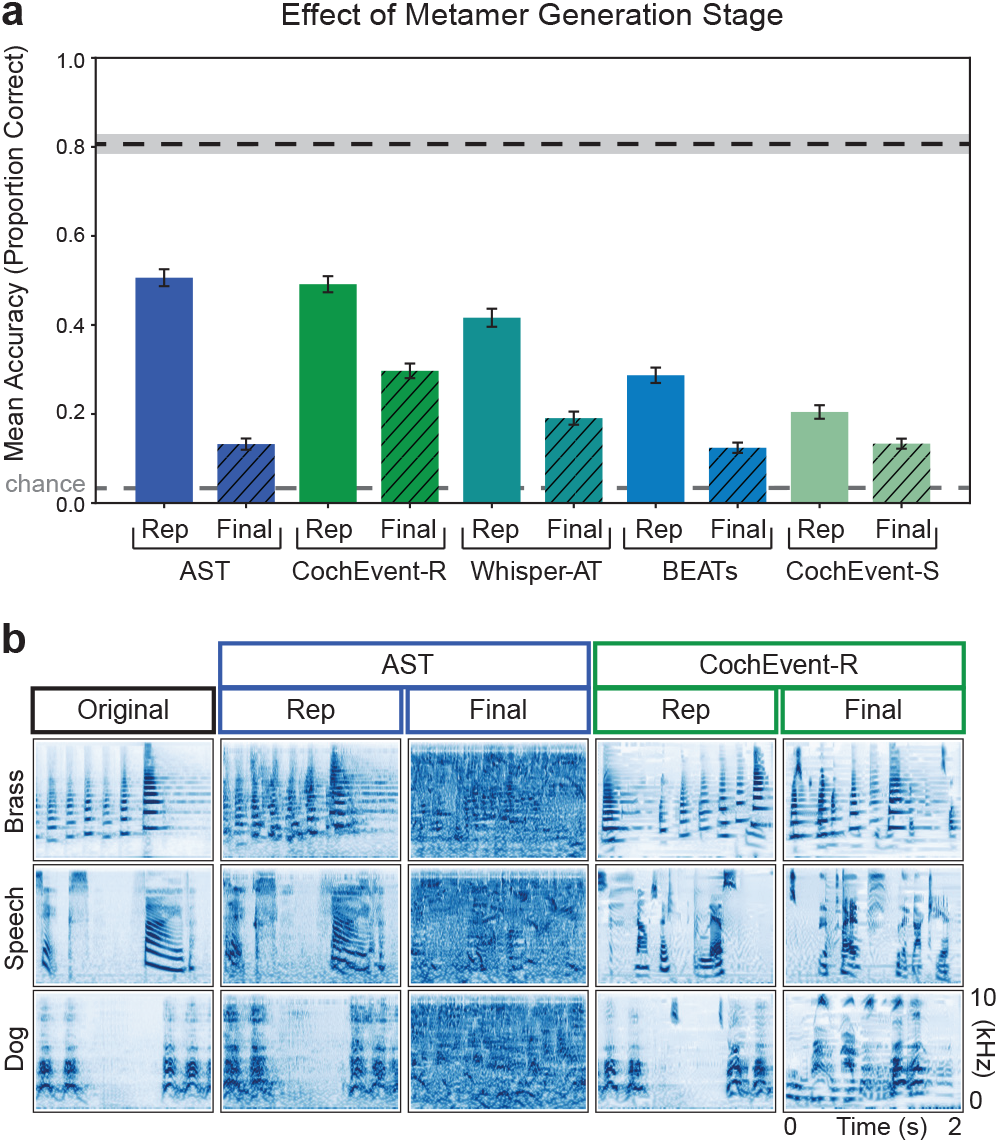
(a) Human recognition of model metamers across audio event recognition models compared to that of natural sounds (*N* = 48). Dotted line denotes human recognition accuracy of natural sounds. Shaded line and error bars denote SEM across participants. Model metamers were generated from the penultimate representation layer (in solid, labeled “Rep”; also plotted in Figure 3) as well as the final classification layer (in hash). (b)Example cochleagrams from “Brass”, “Speech”, and “Dog” categories for AST and CochEvent-R, shown alongside the specific natural sound used as the reference for its synthesis.

**Figure 6:**
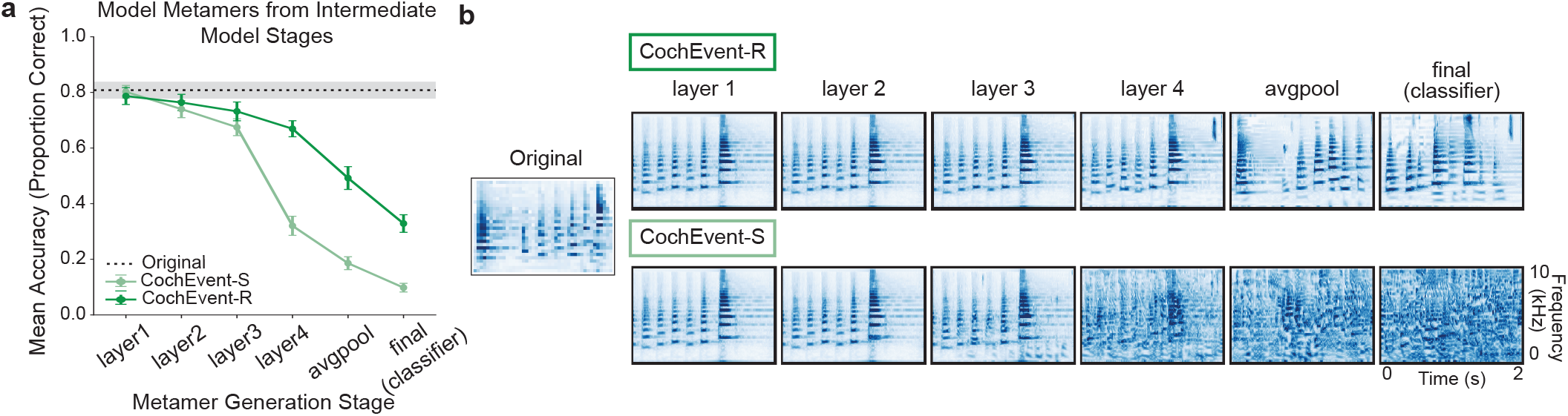
(a) Human recognition of model metamers from selected layers of the CochEvent-S and CochEvent-R model hierarchy (*N* = 17). Early layers (left) are closer to the input representation and generally preserve most of the features necessary for categorization of the sound, while late layers (right) build invariances that support categorization . The dotted line denotes human recognition accuracy of natural sounds. Shaded line and error bars denote SEM across participants. A subset of layers were tested, corresponding to the end of each residual block (layer1, layer2, layer3, layer4), the average pooling layer, and the final classification layer. (b) Example cochleagrams are shown alongside the reference natural sound used as the reference for its synthesis.

## Discussion

DNNs trained on ecologically relevant tasks have emerged as leading candidate models of human representations in multiple sensory domains. In this work, we rigorously tested these models in the domain of auditory event categorization by designing a novel 25-way behavioral benchmark and evaluating the alignment between model and human categorical invariances. We found that both training task and training technique influenced human-model similarities. Specifically, models optimized for auditory event recognition generally produced metamers that were more recognizable to human listeners than those optimized for speech recognition. Furthermore, consistent with previous research in the visual and word-recognition domains, adversarial training significantly improved metamer recognizability. Even though humans can recognize model metamers significantly above chance for some models, a substantial gap remains between overall human classification performance on natural sounds vs. model metamers.

Our results demonstrate that for many models, the optimization objective places constraints on learned categorical invariances. There was a general trend where models optimized for auditory event categorization produced more recognizable model metamers for auditory events than models optimized exclusively in the speech recognition domain. Additionally, for some of the speech recognition models the model metamers for “speech” were correctly categorized as “speech” even though accuracy for the other classes was low (Figure S2). Although the task-specificity of the models may be expected based on the training objective, the field of auditory perception often places a strong emphasis on speech perception benchmarks, which, as demonstrated in our results, may yield models that lack general-purpose capabilities. A notable exception to this trend was Whisper (Radford et al., 2023). Despite being a speech-optimized model, Whisper produced highly recognizable model metamers that rivaled those of top-performing audio event trained models. This is aligned with recent work reporting that Whisper encodes sufficient information about non-speech background sounds, to the extent that adding a readout to Whisper for auditory events can achieve high performance on engineering benchmarks (Gong et al., 2023). While it remains unclear which specific aspects of Whisper’s training or architecture give rise to more human-aligned representations, several key differences relative to the other speech models may be relevant (Supplementary Table S1). In particular, Whisper is trained via a multi-task speech objective on a massive dataset of weakly labeled, real-world audio that includes both speech and background sounds. As noted by Gong et al. (2023), this induces an audio representation that is not noise invariant, but rather achieves robust performance on speech recognition by actively encoding non-speech sounds. This suggests that massive acoustic diversity, combined with multi-task objectives that necessitate preserving background events, may promote a generalized auditory representation. Disentangling the relative contributions of these factors remains an important direction for future work, particularly given that our model selection was intended to broadly sample current architectures and publicly available models rather than test specific hypotheses about model architecture types and training dataset regimes.

Multiple lines of work in both visual and auditory perception have proposed adversarial training as a method to encourage alignment of human and model representations (Dapello et al., 2022; Engstrom et al., 2019b; Feather et al., 2023; Gaziv et al., 2023; Harrington & Deza, 2021). By forcing the models to be robust against small, often human-imperceptible perturbations, adversarial training appears to encourage the development of *perceptually aligned* features for categorization. However, our controlled model comparisons reveal a critical caveat to this hypothesis. While adversarial training improved recognizability for model metamers from the event-trained model (CochEvent-R), it did not have the same effect for the word-trained model (CochWord-R), which produced the least recognizable metamers. This suggests that adversarial training is not a universal solution for human-model alignment; rather, its benefits are most pronounced when the model’s objective is aligned with the human task of interest. This also rules out the possibility that adversarial training is simply encouraging the model to preserve all information (such that model metamers are purely reconstructions of the input). If that were the case, model metamers would be recognizable for both adversarially trained models. Rather, the training process seems to preserve task-relevant features that align with human perception.

Many of the non-adversarially trained models tested here produced model metamers that were recognizable above chance by human observers at the penultimate representation stage, deviating from previous work in vision audio (Feather et al., 2019, 2023). However, even in these models with highly recognizable model metamers, especially AST and Whisper-AT, recognizability substantially degraded at the classification logits. Since metamers effectively reveal aspects of the audio that the model discards, this substantial drop at the final stage reveals that a single linear projection effectively throws away perceptually aligned features to perform the categorization task. In particular, the final classification layer acts as a task-specific decision readout that compresses the representation into a single categorical variable optimized for the training objective (DiCarlo & Cox, 2007), but it potentially prioritizes features that differ from those used by human observers (Geirhos et al., 2020). A rigid single-objective representation fundamentally diverges from hypotheses about how biological systems make decisions (Gold & Shadlen, 2007), and suggests that future models might incorporate robust representations supporting multiple tasks (Johnston & Fusi, 2023) or aspects of mixed selectivity (Fusi et al., 2016). This additionally underscores the problem of using accuracy alone for model-human comparison: high accuracy can be achieved by discarding information that is critical for human categorization. Crucially, the high recognizability of metamers at earlier representation stages provides evidence that the performance gap at the final decision stage reflects learned invariances of the model rather than fundamental limitations of the metamer synthesis framework. Given that we see improvement with some types of training procedures (e.g. adversarial training) we anticipate that future architectures or training objectives can further close this categorization gap through the final classification stage.

Finally, the design of our 25-way behavioral paradigm provides a rigorously controlled, ecological benchmark for human-model evaluations. Constructing a task for auditory event perception is non-trivial; unlike in visual recognition where the definition of an “object” tends to be well agreed upon, the precise definition of an “auditory object” remains opaque. While we motivated our choice of auditory categories based on available datasets and labeling schemes, the human-model comparisons might differ if we examined the categorization at different scales. To navigate this, we have attempted to build some of these into the categorization choices (for instance, by including multiple “music” and “animal” categories), and we demonstrate that our results are consistent if we analyze human and model responses based on the five superordinate categories (Figure S1), but the precise choice of categories may still result in differences between human and model evaluations, which could be explored in future work. For a given model, there is also structured variability in metamer recognizability across different auditory categories (Figure S2). This variability within a model could be used to identify if the model has particularly inhuman invariances for certain types of sounds (for instance, ones with certain types of spectrotemporal modulation patterns). Further, natural sounds often exhibit acoustic information that easily blurs between distinct semantic categories. Our benchmark attempts to navigate this complexity by incorporating many examples of each category to reveal systematic confusions in auditory categorization patterns for natural sounds presented in isolation. For example, we observed a systematic human confusion between “cat” and “car”, which could be attributed to samples of a low-frequency cat purr. These error patterns are not failures, but reveal signatures of human-like processing that are desirable in a model of perception. By formalizing these comparisons with an engaging task that yields high-quality behavioral data, our work establishes the foundation for building and evaluating future models of auditory event categorization. While identifying the exact mechanistic computations underlying human auditory categorization remains an open challenge, future work can utilize our benchmark to systematically probe the underlying representations and build models that better align with human perception.

## Acknowledgments

We thank Maggie Henderson and Ross Williamson for helpful feedback throughout this project and Audra Irvine and Jenah Black for helping with the online data collection process. We also thank the NeuroAI SuperLab at CMU for comments on this work. A.N.’s effort was supported by the Office of Naval Research (N00014-53618-1-2069).

## Supplemental Material

### SM.1: CochEvent-R Model Training

The adversarially trained CochEvent-R model was optimized on the Auditory Event component of the WordSpeaker-Noise (WSN) dataset (Feather et al., 2019). The model had a ResNet50 (He et al., 2016) backbone with a cochleagram front end, and training procedures were matched to those reported for the CochEvent-S model (Tuckute et al., 2023). At each step of training, an adversarial example was generated for the cochleagram representation for each example in the training batch, such that the example had maximum *l*_2_-norm = 1. The adversarial examples were untargeted (generated by maximizing the training loss), and were generated with a step size of 0.5 and five steps of projected gradient descent, using a modified version of the robustness library (Engstrom et al., 2019a) that had additions for training audio models (Feather et al., 2023). The model was trained on the 517way audio event recognition task consisting of a subset of AudioSet data using a Binary Cross Entropy Loss. The model was trained for 48 epochs of the WSN AudioSet training data with a batch size of 256, gradient clipping (max *l*_2_-norm = 1.0), initial learning rate of 0.001, and a warm-up learning rate for the first 500 batches (learning rate = <initial learning rate> / (500-i) where i is the batch number).

### SM.2 Additional details for human behavior experiments

All experiments with human participants were approved by the Institutional Review Board at Carnegie Mellon University and conducted with the informed consent of the participants. Experiments were created and hosted using the Gorilla Experiment Builder (www.gorilla.sc; Anwyl-Irvine et al., 2020) and were conducted using the Prolific platform (www.prolific.com). Inclusion criteria on prolific consisted of location based in the United States, English as first language, and self-reported no hearing difficulties. Each session took approximately 30 minutes to complete and participants were paid$6 for completion of the experiment. For reproducibility, each experiment is hosted on Gorilla Open Materials:

(1)natural sounds experiment (Figures 1,2)

https://app.gorilla.sc/openmaterials/1297636

(2) main model metamer experiment (Figures 3,4)

https://app.gorilla.sc/openmaterials/1297642

(3) all layers model metamer experiment (Figure 6)

https://app.gorilla.sc/openmaterials/1297690

Before entering the full behavioral task, participants completed a “headphone check” (Milne et al., 2021), to ensure a high-quality audio environment. Participants who failed the headphone check had to return the experiment and a new participant was recruited (18 failures total across all experiments). Experimental data were only included in the analysis if participants passed at least 10 of 11 attention checks. During these attention checks, participants were verbally instructed to click on a specific category icon. The verbal instructions were generated by Voicemaker (https://voicemaker.in/). For the experiments included in this paper, no participants failed the attention screening check.

To help participants familiarize themselves with the 25 categories, we included an introduction phase in which we presented an example sound for each category that was not part of the main set of 275 stimuli. Participants could replay the sound multiple times during this training phase. At the end of the training phase, participants completed three practice trials consisting of stimuli that were not included in the test trials, and participants had to get at least two out of three of these trials correct to proceed to the full experiment.

### SM.3 Data and code availability

To ensure reproducibility and make our benchmark accessible to the community, all stimulus sets, behavioral data, and code for metamer generation and model evaluations are publicly available via a GitHub repository: https://github.com/Feather-Lab/env-sound-metamers.

In *Proceedings of the 9th Conference on Cognitive Computational Neuroscience*, New York, NY, USA, 2026.

In *Proceedings of the 9th Conference on Cognitive Computational Neuroscience*, New York, NY, USA, 2026. doi:10.32470/2gj54og Copyright 2026 by the author(s). Licensed under CC BY 4.0.

**Table S1:** Comparison of model architectures, training objectives, and datasets. Models differ in scale, supervision, and the diversity of training data, which may contribute to differences in alignment with human perception. SSL stands for self-supervised learning. CE stands for cross-entropy.

| Model | Architecture | Objective / Loss | Optimization | # Param | Training Data |
| --- | --- | --- | --- | --- | --- |
| wav2vec2.0-Large | CNN + Transformer encoder | Contrastive SSL | Adam | 317M | 960 hours, primarily speech |
| HuBERT-Large | CNN + Transformer encoder | Masked prediction | Adam | 317M | 960 hours, primarily speech |
| AST | Transformer (ViT) | CE | Adam | 86M | ~2M clips, natural sounds |
| BEATs | Transformer (SSL + finetuning) | SSL + CE | Adam | 90M | ~2M clips, natural sounds |
| Whisper-Large V1 | Encoder-Decoder Transformer | CE (Multitask ASR) | Adam | 1550M | ~680k hours, web audio of speech |
| Whisper-AT | Whisper encoder + Transformer | CE | Adam | 1550M | ~2M clips, natural sounds |
| CochWord-R | Cochleagram + ResNet-50 | CE | SGD w/ momentum | 26M | ~0.23M speech clips, speech superimposed with natural sounds |
| CochEvent-S | Cochleagram + ResNet-50 | CE | SGD w/ momentum | 26M | ~0.72M natural sound clips, natural sounds superimposed with speech |
| CochEvent-R | Cochleagram + ResNet-50 | CE | SGD w/ momentum | 26M | ~0.72M natural sound clips, natural sounds superimposed with speech |

**Table S2:** Summary of model architectures and specific representation stages utilized for metamer generation. ASR refers to automatic speech recognition. LS, AS, and IN respectively refer to LibriSpeech, AudioSet, and ImageNet. Whisper* denotes a frozen encoder backbone where only the top-level classification head (TL-Tr) was trained on AudioSet. Word-Speaker-Noise dataset comes from Feather et al., 2019. All models performing audio event classification were trained to predict AudioSet labels.

| Model | Training Dataset | Training Task | Metamer Generation Layer |
| --- | --- | --- | --- |
| **Audio Event Recognition Models** |  |  |  |
| AST [Gong et al., 2021] | IN + AS (2M clips) | Audio Event Classif. | Encoder Output (L12), Logits |
| BEATs-Iter3+ [Chen et al., 2022] | AS (2M clips) | Audio Event Classif. | Encoder Output (L12), Logits |
| Whisper-AT [Gong et al., 2023] | Whisper* + AS | Audio Event Classif. | TL-Tr Output, Logits |
| CochEvent-S [Tuckute et al., 2023] | Word-Speaker-Noise | Audio Event Classif. | Avgpool, Logits |
| CochEvent-R | Word-Speaker-Noise | Audio Event Classif. | Avgpool, Logits |
| **Speech Recognition Models** |  |  |  |
| wav2vec 2.0-Large [Baevski et al., 2020] | LS (960h) | ASR (CTC) | Encoder Output (L24) |
| HuBERT-Large [Hsu et al., 2021] | LS (960h) | ASR (CTC) | Encoder Output (L24) |
| Whisper-Large V1 [Radford et al., 2023] | Multilingual (680k h) | Multi-task ASR | Encoder Output (L32) |
| CochWord-R [Feather et al., 2023] | Word-Speaker-Noise | Word Recognition | Avgpool |

**Table S3:** We report the number of metamers generated, feature dimensionality, and optimization performance (Spearman ρ) for each model layer. Optimization hyperparameters are abbreviated as follows: I: number of iterations per repeat; N: number of repeats (*I × N* = total number of steps taken to generate metamers); Z: initial maximum *L*_2_ norm (η) for gradient descent; SRD: step size reduction factor (the current η is divided by *SRD* after every *I* iterations); E: noise scale used for initialization (e.g., E-5 denotes 10^*−*5^). For example, “I3000 N8 Z1.0 SRD2.0 E-5” indicates 8 repeats of 3,000 iterations with an initial η of 1.0, halved every 3,000 steps.

| Model Metamer Generation Layer | Number Metamers | Number Features | Optimization Hyperparameters | Median Spearman $\rho$ at Layer | Median Spearman $\rho$ at Final Layer |
| --- | --- | --- | --- | --- | --- |
| wav2vec2 |  |  |  |  |  |
| Encoder L24 | 267 | 101376 | I3000 N8 Z1.0 SRD2.0 E-5 | 0.99669 | 0.99958 |
| HuBERT |  |  |  |  |  |
| Encoder L24 | 274 | 101376 | I3000 N8 Z1.0 SRD2.0 E-5 | 0.99947 | 0.99985 |
| AST |  |  |  |  |  |
| Encoder L12 | 266 | 932352 | I3000 N20 Z1.0 SRD2.0 E-5 | 0.98801 | 0.99399 |
| Logits | 274 | 527 | I500 N25 Z2.0 SRD1.50 E-5 | 0.99989 | 0.99989 |
| BEATs |  |  |  |  |  |
| Encoder L12 | 275 | 73728 | I1000 N15 Z2.0 SRD1.7 E-5 | 0.99899 | 0.99995 |
| Logits | 273 | 527 | I3000 N9 Z2.0 SRD2.0 E-7 | 0.99980 | 0.99980 |
| Whisper |  |  |  |  |  |
| Encoder L32 | 274 | 1920000 | I3000 N8 Z1.0 SRD2.0 E-5 | 0.99858 | 0.99998 |
| Whisper-AT |  |  |  |  |  |
| TL-TR Out | 269 | 16384 | I1000 N10 Z1.3 SRD2.0 E-5 | 0.99669 | 0.99958 |
| Logits | 275 | 527 | I1000 N10 Z1.3 SRD2.0 E-5 | 0.99940 | 0.99940 |
| CochWord-R |  |  |  |  |  |
| Avgpool | 274 | 2048 | I3000 N8 Z1.0 SRD2.0 E-7 | 0.99319 | 0.99992 |
| CochEvent-S |  |  |  |  |  |
| Avgpool | 273 | 2048 | I3000 N20 Z2.0 SRD1.5 E-7 | 0.99574 | 0.99998 |
| Logits | 270 | 517 | I3000 N20 Z2.0 SRD1.5 E-7 | 0.99981 | 0.99981 |
| CochEvent-R |  |  |  |  |  |
| Avgpool | 274 | 2048 | I3000 N15 Z1.3 SRD2.0 E-5 | 0.99835 | 0.99997 |
| Logits | 270 | 517 | I3000 N15 Z1.3 SRD2.0 E-5 | 0.99985 | 0.99985 |

**Figure S1:**
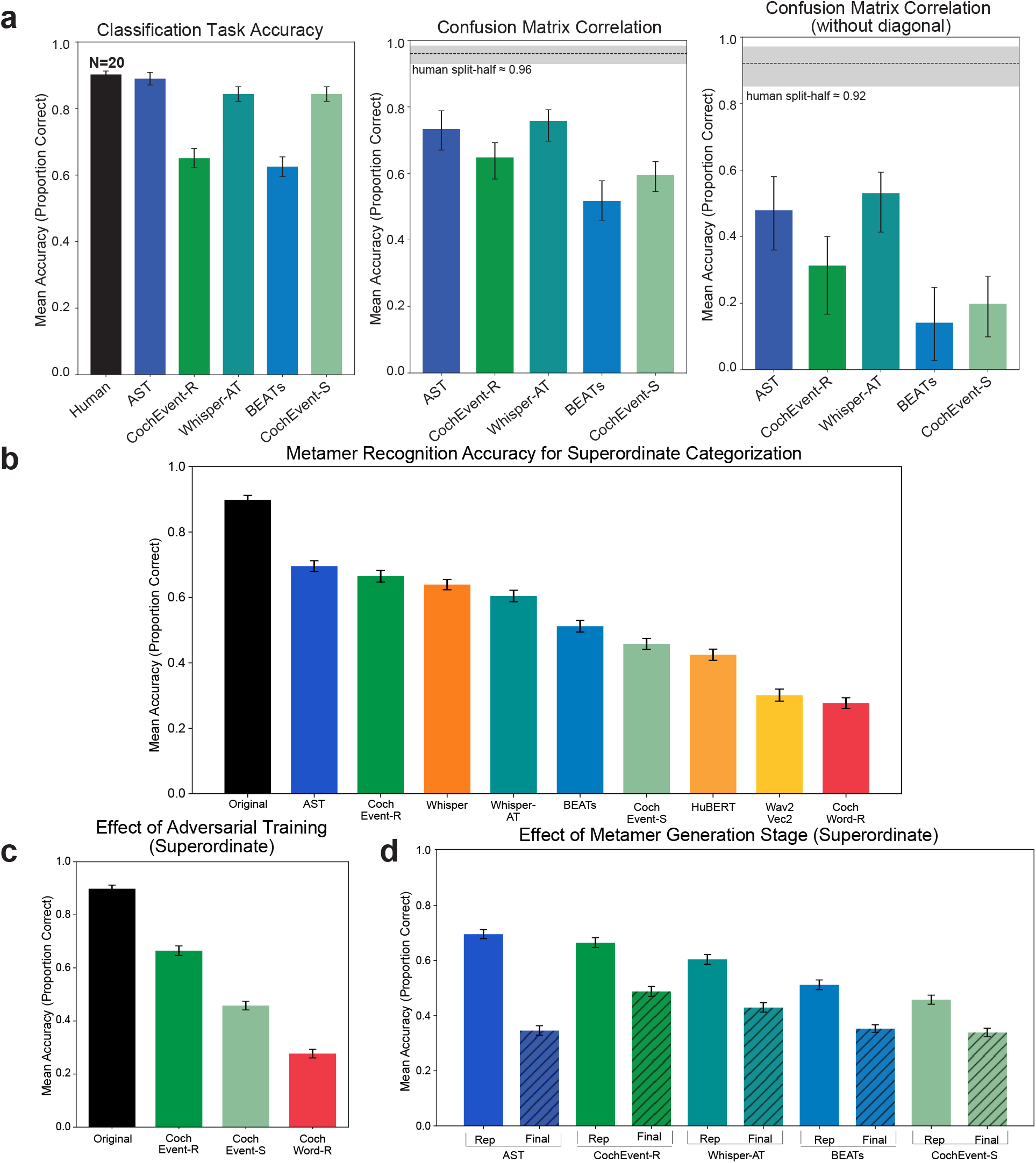
(a) Superordinate-category version of Figure 2. We replicated the analysis in Figure 2 after collapsing the original 25 fine-grained categories into 5 superordinate classes (Human sounds, Animal, Music, Natural sounds, Sounds of things; see Figure 1b). Confusion matrices were constructed using these superordinate labels, such that responses were considered correct if the predicted and true labels matched at the superordinate level. Dashed line indicates human split-half reliability (Spearman-Brown corrected). Shaded lines and error bars follow same procedures as Figure 2. (b & c & d) Superordinate-category version of Figures 3b, 4, 5. Accuracy of model metamer experiment computed at superordinate level. Error bars denote SEM across participants.

**Figure S2:**
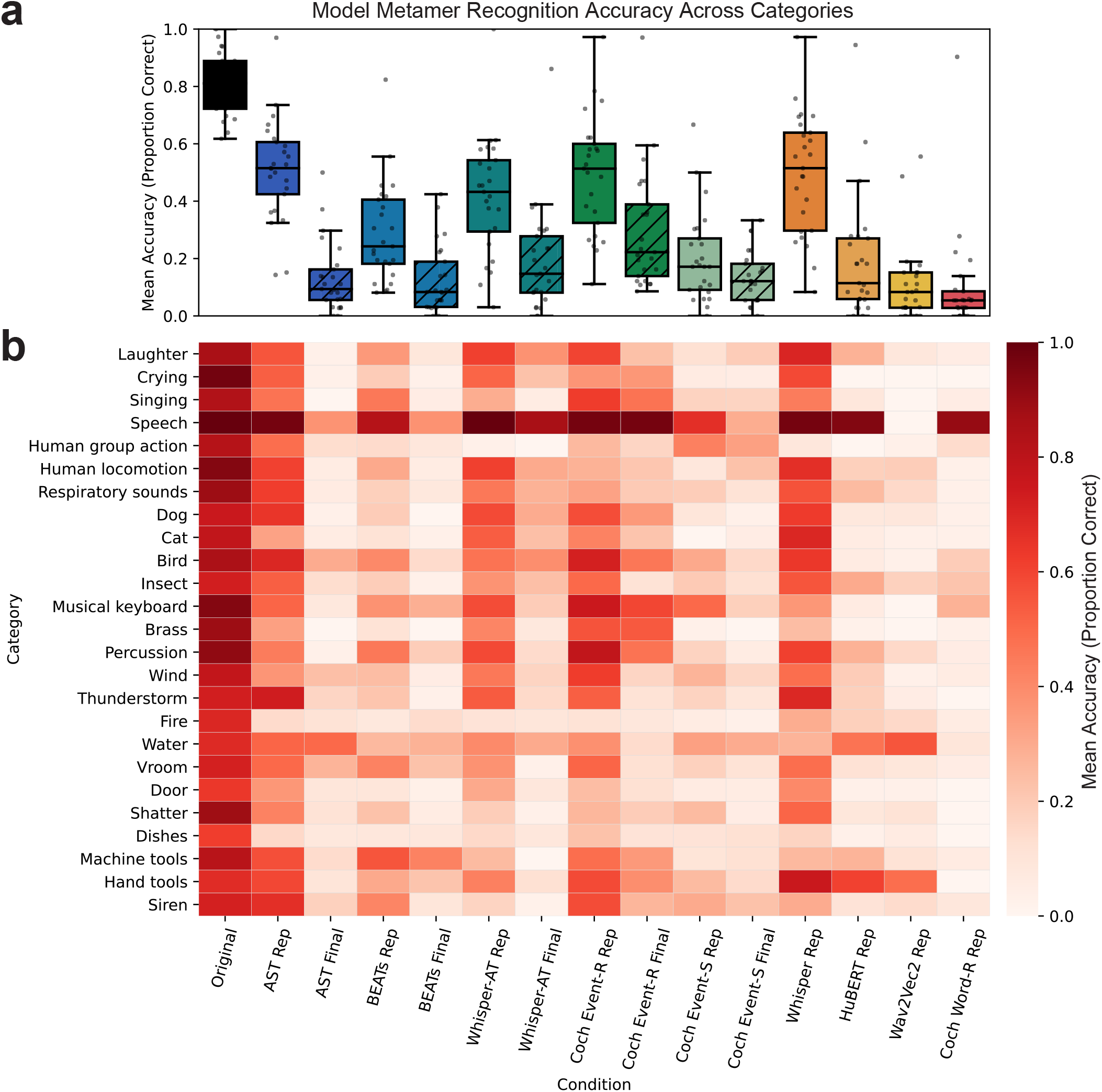
Replotted metamer recognition accuracy for human participants across all model conditions across the 25 different category conditions. (a) Category performance as a box and whisker plot with individual dots correspond to each category. (b) Heatmap showing recognition accuracy for each category (rows) and model condition (columns).

**Table S4:**
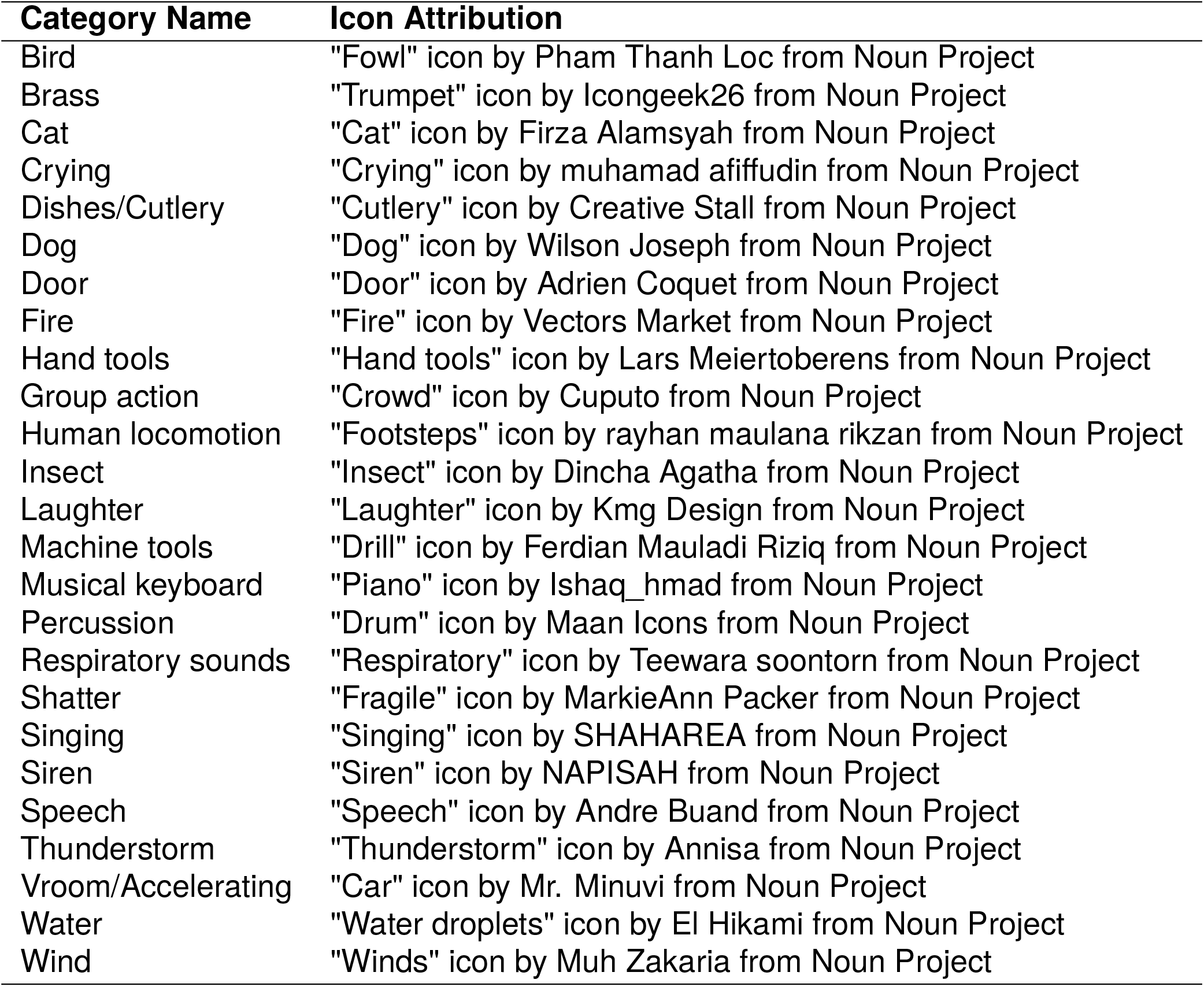
Icon Attributions.

